# Direct monitoring of unfolded nascent polypeptides by the CCR4-NOT complex contributes to protein quality control through selective mRNA deadenylation and decay

**DOI:** 10.64898/2026.10.08.757698

**Authors:** Guillaume Caulier, Fabienne Mauxion, Bertrand Seraphin

## Abstract

Accumulation of aberrant and unfolded proteins induce cellular stresses and contributes in human to diseases and aging. Several cellular pathways combat the formation of these deleterious molecules. Recently, processes limiting the production of unwanted polypeptides through the co-translational degradation of their messenger RNAs (mRNAs) have been identified but remain poorly understood. In *Saccharomyces cerevisiae*, Caf130 was proposed to regulate the levels of mRNAs encoding some ribosomal proteins, principally Rpl4, when they do not fold properly. We report the functional and structural characterization this process. Biochemical purifications, interaction assays, and in vivo analyses combined with Alphafold modelling define the organization and function of the Caf130 module of the CCR4-NOT complex and associated factors that mediate this process. Our data reveal the critical role of different forms of the Nascent protein Associated Complex (NAC) to dock the CCR4-NOT complex onto translating ribosomes. The CCR4-NOT complex directly recognizes some nascent unfolded proteins emerging from the ribosome tunnel triggering deadenylation of the mRNAs encoding them. Our data further reveal unsuspected evolutionary and functional similarities between yeast and mammalian proteins involved in the quality control of nascent proteins through mRNA degradation. Altogether, our data indicate the existence of a conserved machinery contributing to protein quality control by triggering the decay of mRNAs encoding deleterious polypeptides in eukaryotes.

## Introduction

The accurate regulation of mRNA stability is an essential process that contributes to maintaining protein homeostasis, particularly in response to cellular stress and fluctuations in gene expression demand. In eukaryotes, the major mRNA decay pathway is initiated by deadenylation of the poly(A) tail, primarily mediated by the CCR4-NOT complex, a multi-subunit deadenylase conserved across species (Wahle & Winkler, 2013; Collart, 2016; Caulier et al., 2025). This evolutionarily conserved complex integrates multiple features present in mRNAs to control their deadenylation that subsequently induce their decay (Caulier et al., 2025). The CCR4-NOT assembly is composed of eight core components in *Saccharomyces cerevisiae* (Denis, 1984; Sakai et al., 1992; Liu et al., 1998; Oberholzer & Collart, 1998; Caulier et al., 2025) with the Not1 protein serving as a central scaffold that organizes the complex. Deadenylation is mediated by the Ccr4 subunit (Basquin et al., 2012; Bhaskar et al., 2013, 2015; Caulier et al., 2025). All other subunits of the yeast CCR4-NOT complex (Caf1/Pop2, Caf40, Not2, Not3/Not5, Not4, Caf130) are conserved in its human counterpart except for Caf130. Reciprocally, the CNOT10 and CNOT11 subunits that docks on the N-terminus of CNOT1 in the human assembly have no known homologues in yeast (Mauxion 2013) while CNOT4 interacts only transiently with the complex in metazoan. Beside the CCR4 deadenylase, the other complex components have been implicated in complex integrity, ubiquitination, and/or targeting of the complex to substrate mRNAs.

The CCR4-NOT complex function has recently emerged as being tightly connected to translation. A cryo-electron microscopy study in yeast revealed that Not5 interacts with ribosomes stalled at rare codons, recognizing specific ribosomal conformations of its E- and A-sites. This induces the deadenylation and degradation targeting the associated mRNAs (Buschauer et al., 2020) consistent with codon biases being a major parameter determining mRNA turnover rates. A similar mechanism has been identified in human cells, where the homologous CNOT3 factor recognizes specific ribosomal features at the E-site as well as the P-site tRNA to promote decay of inefficiently translated transcripts, underscoring the evolutionary conservation of this pathway (Absmeier et al., 2023; Zhu *et al.,* 2024).

Interactomic studies have revealed that the Caf130 subunit of the yeast CCR4-Not complex connects with Btt1 a subunit of the Nascent polypeptide-Associated Complex (NAC) (Ito et al., 2001; Krogan et al., 2006; Cui et al., 2008). NAC, a dimeric complex present in archaea and eukaryotes, binds ribosome in the vicinity of the ribosome tunnel from which emerge nascent polypeptides. NAC associates with additional factors such as aminopeptidase and acetylase to fold and mature newly synthesized proteins. While archaeal NAC are homodimeric, their eukaryotic counterparts are heterodimeric being formed of two related alpha- and beta-subunits. In the yeast *Saccharomyces cerevisiae* NAC, the Egd2 NAC alpha-subunit interacts with either one of two alternative beta-subunits Egd1 or Btt1 to form NAC^Egd1^ or NAC^Btt1^, respectively. Interestingly, NAC^Egd1^ represents the majority of the NAC complex in yeast cells as its abundance is estimated to be 20-50 fold higher than NAC^Btt1^.

Based on the suppression of the slow growth of cells lacking Acl4, a dedicated chaperone binding the Rpl4 ribosomal protein and facilitating its incorporation in nascent ribosomes, Pillet et al. (2022) proposed a model implicating Caf130, Btt1 and Cal4 bound to the N-terminus of Not1 to recognize excess nascent Rpl4 and trigger deadenylation and degradation of supernumerary Rpl4 mRNA. These preluding observations did not elucidate the precise function and structural organization of the Caf130 module within the CCR4-NOT complex nor define how the nascent Rpl4 protein would be recognized.

**Here**, we investigate the architecture and role of the Caf130 module of the yeast CC4-NOT complex. By combining biochemical purifications, protein interaction assays, *in vivo* functional analysis and structural modelling using AlphaFold, we define the organization of this module. Functional and genetic assays further demonstrate that the complex indeed controls deadenylation of the Rpl4 mRNA and reveals further that involves a full NAC complex, NAC^Egd1^ or NAC^Btt1^ participating to this process to different extent. We also provide evidence that NAC acts as key adaptor to dock the CCR4-NOT complex onto translating ribosomes where it is directly involved in recognition of the nascent Rpl4 protein. Additionally, our data identify an unanticipated functional and structural similarity between the yeast protein Caf130 and mammalian CNOT11. Overall, our data reveal that the CCR4-NOT complex directly monitors translating ribosomes to eliminate mRNAs encoding deleterious nascent polypeptides. This indicates the existence of a conserved machinery that participates to protein quality control and limits accumulation of potentially harmful proteins in eukaryotes, thereby contributing to combat proteinopathies in human.

## RESULTS

### Interactome of the N-terminal module of the CCR4-NOT complex in Saccharomyces cerevisiae

To get insights into the organization and cellular integration of the N-terminal module of CCR4-NOT in *S. cerevisiae*, we purified Caf130 using the TAP purification method (Puig et al. 2001). Plotting the level of the copurifying partners estimated by mass spectrometry versus their abundance in extract (Figure 1A) revealed the enrichment of the CCR4-NOT complex, including the recently described Cal4 associated factor as the top partners. Additional co-purifying factors included the two NAC α and β subunits Egd2 and Btt1. The presence of Egd2 was unexpected as previous analyses had suggested that Btt1 interacted with Caf130 in its absence (Pillet et al. 2022). Interestingly, the alternative yeast β-NAC subunit, Egd1, was among the less enriched partners despite being more abundant than its Btt1 counterpart demonstrating that association with the latter is specific. Other complexes and proteins associated with Caf130 included the Pbp1 complex that has been functionally connected to mRNA decay and translation, and some translation factors including ribosomal proteins (Figure 1A). Concerning the latter, one should take into account that, given their very high abundance in extracts, that largely exceeds the level of Caf130, one can expect at most a modest enrichment.

**Figure 1:**
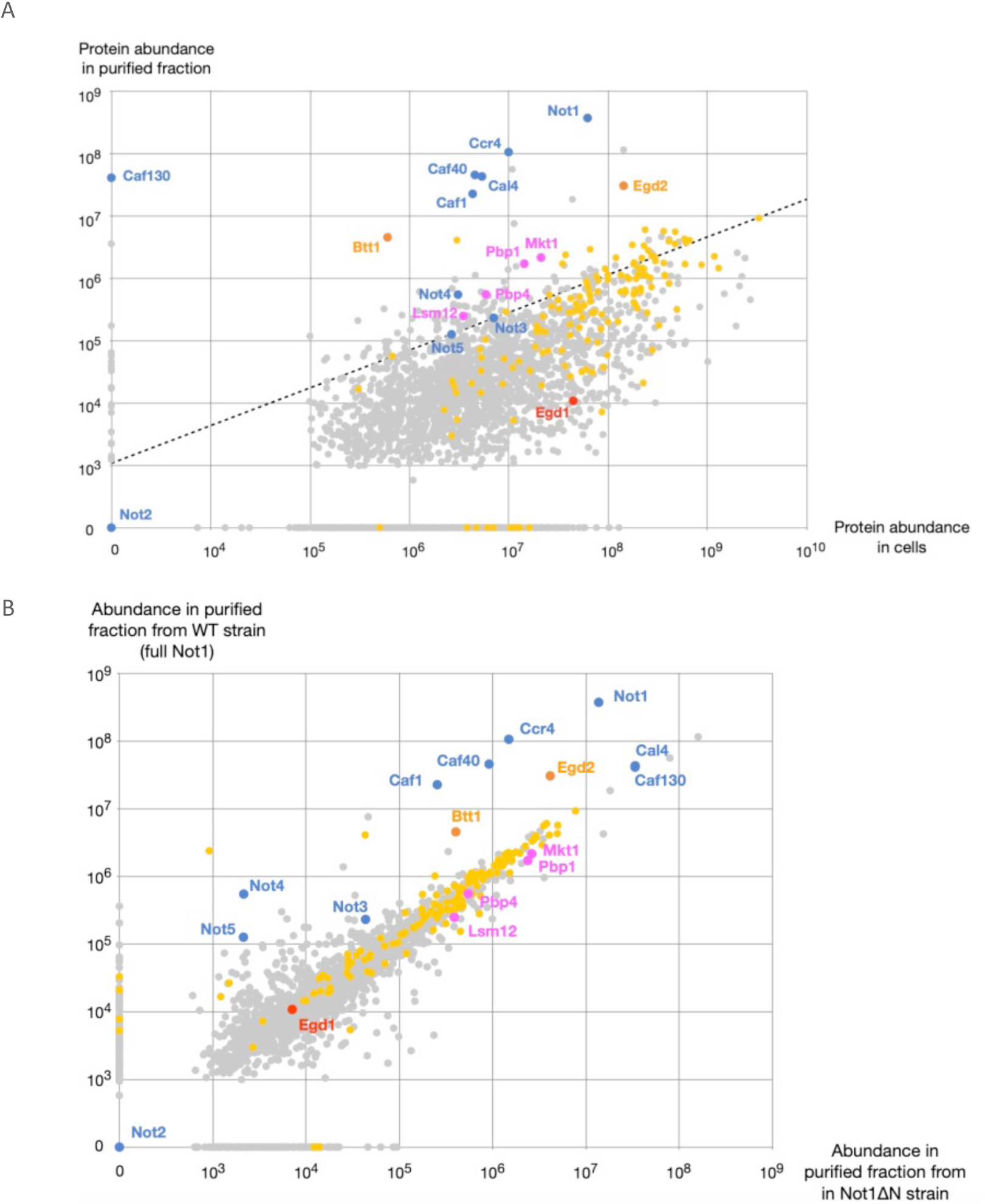
Identifying partners of Caf130 by TAP purification – mass spectrometry. (A) Proteins were identified and quantified by label-free mass spectrometry following TAP purification of Caf-130-TAP and from the starting extract. The log of the protein abundance in the TAP-purified sample is plotted as a function of the log of the protein abundance in extract based on the extracted ion chromatography intensity of the peptides. Proteins not identified in the TAP-purified sample or in extract are presented on the axes labelled “0”. Points above the dotted line represent the 5% most enriched proteins taking into account that background signal increases with protein abundance in extract. For this purpose, the dotted line is parallel to the best fit line of the whole distribution excluding points identified only in one of the two samples. Points are colored according to the protein function: blue: CCR4-NOT complex including Cal4, orange: NAC^Btt1^, red: Egd1, pink: Pbp1 complex, yellow: ribosome and translation factors, grey: all other points. (B) Comparison of the proteins identified and quantified following the TAP purification of Caf-130-TAP presented in (A) with those from the TAP purification of Caf-130-TAP in a Not1 (Δ*N-terminal*) strain. The log of the protein abundance in the TAP-purified samples are plotted with proteins not identified in one or the other sample presented on the axes labelled “0”. Color are as in (A).

We and others have shown that Caf130 associate with the N-terminus of Not1 (Pillet et al. 2022). Thus, to assess the specificity of the purification, we performed a second TAP purification of Caf130 from a strain carrying a deletion of the 162 N-terminal residues of Not1. Comparing the two purifications (Figure 1B) demonstrated that association of Caf130 with other CCR4-NOT subunits occurred via its interaction with Not1. The only exception was Cal4 that was as efficiently co-purified from the two strains indicating a direct interaction with Caf130. Interestingly, Egd2 and Btt1 were co-purified from both strains but slightly more efficiently from the strain carrying full-length Not1. This suggests that the NAC^Btt1^ complex binds Caf130-Cal4 but that this interaction is stabilized when the N-terminus of Not1 is present. Other co-purifying factors including the Pbp1 complex and translation factors were enriched to a similar extent whether the N-terminus of Not1 was present or not.

Overall, these data suggest that Caf130 associates with Cal4, Egd2, Btt1, and the N-terminus of Not1 to form a specific assembly that does not appear to contain additional component.

### Validating and mapping Caf130 interaction with partners

To validate the affinity purification – mass spectrometry results and decipher the network of interaction between the components of the Caf130 module, we employed the two-hybrid tests. The host yeast strain was transformed with plasmids encoding full-length Caf130 fused to the Gal4 DNA Binding Domain (DBD) together with Not1, Egd1, Btt1, Egd2, or Cal4 fused to the Gal4 transcription Activation Domain (AD). Empty vectors were also included as specificity controls. Interaction signals were monitored by quantifying β-galactosidase production. Significant signals were detected for Caf130 with Btt1, Not1, or Cal4 (Figure 2A). This supports an interaction between these factors. Surprisingly, no signal was detected for the Egd2 - Caf130 pair. However, negative results are poorly informative because a given fusion protein may poorly enter the nucleus or interfere with transcription activation. Further, high level of endogenous Egd1 and Egd2 proteins may compete with the corresponding fusion proteins, while the low-copy Btt1 factor may not. Overall, the two-hybrid assay supports interactions between Caf130 and Btt1, Cal4 or Not1.

**Figure 2:**
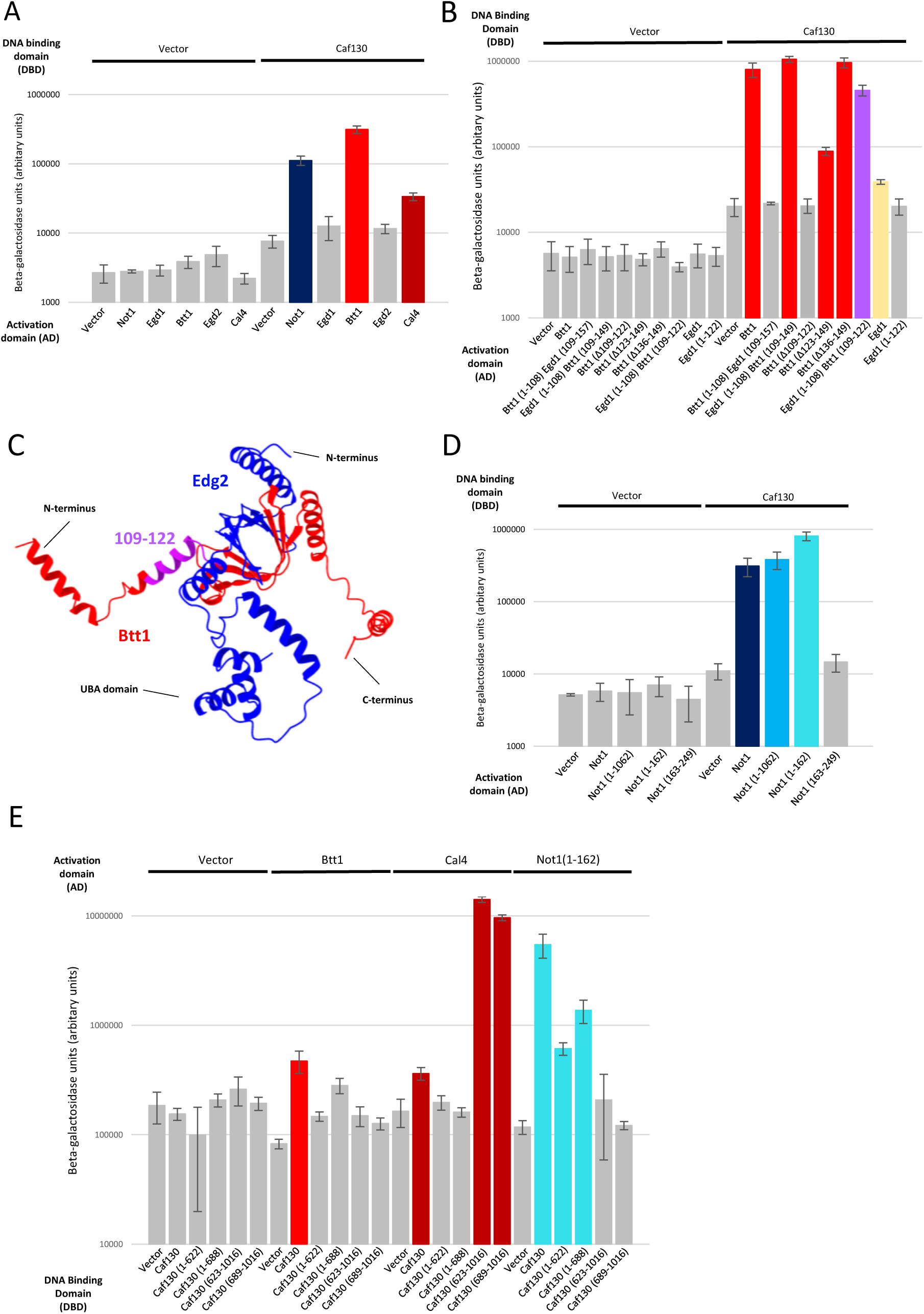
Validating and mapping interactions of Caf130 with its partners. (A) Results of interaction tests between Caf130 and Not1, Egd1, Btt1, Egd2 and Cal4. Protein interactions were monitored by quantitating β-galactosidase activity (arbitrary units) in the two-hybrid assay. Three biological replicates were tested and average and standard error values were plotted on a logarithmic scale. A Kruskal-Wallis test rejected homogeneity with a P value lower than 0.0001. Signals corresponding to background are represented in grey. Positive results were observed for Not1 (dark blue), for Btt1 (red) and for Cal4 (dark red). (B) Mapping of the region of Btt1 required for interaction with Caf130 using he two-hybrid assay. Conditions were the same as in (A). A Kruskal-Wallis test rejected homogeneity with a P value lower than 0,01. Positive signals were detected for Btt1 (red), for Egd1(1-108)-Btt1(109-149) (red), for Btt1(Δ123-149) (red), for Btt1(Δ136-149) (red), for Egd1(1-108)-Btt1(109-122) (purple), and a slight positive signal for Egd1 (yellow). Note that background absolute levels and signals (arbitrary units) varied from one experiment to the next but that relative intensities were conserved. (C) The region required for Btt1 interaction with Caf130 (amino acids 109-122, purple) is plotted on an AlphaFold3 prediction of the *S. cerevisiae* NAC^Btt1^ structure complex with other residues of Btt1 in red and Egd2 in blue. (D) Mapping the region of Not1 required and sufficient for interaction with Caf130 using he two-hybrid assay. Conditions were the same as in (A). A Kruskal-Wallis test rejected homogeneity with a P value lower than 0.0001. Positive signals are shown for Not1 (dark blue), for Not1 (1-1062) (light blue) and for Not1 (1-162) (cyan). (E) Egd2 is important for interaction between Caf130 and Btt1. Conditions were the same as in (A), except that the assays were carried out in a Δ*egd2* yeast host strain. For Caf130 and Btt1, a weak positive result was observed (red). For Cal4, positive results were observed for Caf130, Caf130(623-1016), Caf130(689-1016) (dark red). For Not1(1-162), positive results were observed for Caf130, Caf130(1-622), Caf130 (1-688) (cyan).

To assess whether Egd2 is important implicated in the Caf130 module assembly, we generated an Egd2 deletion in the yeast host strain used to perform two-hybrid assays. This host strain was transformed with plasmids encoding full-length Caf130 and truncation variants fused to the Gal4 DNA Binding Domain (DBD), together with Btt1, Cal4, or Not1(1-162) fused to the Gal4 AD. Empty vectors were included as specificity controls. In the absence of Egd2, the signal for the Caf130 - Btt1 pair is significantly reduced, whereas signals corresponding to Caf130 interaction with Cal4 and Not1 remain unchanged (Figure 2E). This argues that Egd2 is required for efficient interaction between Caf130 and Btt1, supporting its presence in the Caf130 module.

The results obtained above suggest that the two-hybrid assay could map residues differing between Btt1 and Egd1 that are responsible for their differential binding to Caf130. Indeed, Btt1 and Egd1 have the same structural organization and share 46% identity, but behave in an opposite manner in this interaction assay. Truncated and chimera of Btt1 and Egd1 proteins were constructed to map their interactions with Caf130 using two-hybrid assays (Figure 2B-S2). Those confirmed interaction between Btt1 and Caf130 and showed only background-level or a slight interaction between Caf130 and Egd1, consistent with earlier findings (Figure 2A-S2). Since Btt1 and Egd1 differ mainly in their C-terminus (Figure S1), we tested whether these residues were implicated in binding specificity. When the Btt1 region (109-149) was substituted by the homologous Egd1 region (109-157), the resulting protein could no longer interact with Caf130 (Figure 2B-S2). Conversely, grafting the Btt1 region (109-149) onto Egd1(1-108) enabled the chimera to bind Caf130 (Figure 2B-S2). This indicates that Btt1 C-terminal region confers the ability to bind Caf130 when present in a β-NAC subunit. Further deletion of amino acids 109-122 abolished binding, indicating that this region is critical for interaction with Caf130. To confirm this observation, a chimera of Egd1 (1-108) with Btt1(109-122) was constructed and showed strong signal for binding to Caf130, whereas the truncated Egd1 (1-122) control did not. Overall, these results demonstrate that, in the context of a β-NAC scaffold, amino acids 109-122 of Btt1 are necessary and sufficient for interaction with Caf130, while downstream residues are not necessary. This amino acid region corresponds essentially to a α helix in a Btt1 structural model (Figure 2C).

To determine the region of Not1 necessary for Caf130 binding (Figure 2D), a series of Not1 truncations were analysed by two-hybrid assays. Given that Not1 and its derivatives tend to induce transcription when bound to the DBD, a single orientation was tested. Positive signals were observed with Caf130 in the presence of both full-length Not1 and derivatives encompassing the Not1 N-terminus. The smallest region of Not1 interacting with Caf130 contained amino acids 1-162. Consistently, constructs lacking this region, e.g., Not1(163-249), failed to interact with Caf130. Consistent with previous data, our results indicate that Caf130 specifically binds to the extreme N-terminal region 1-162 of Not1. Notably, this Not1 region is sufficient for the interaction and is significantly conserved in Eukaryotes (Figure S3).

Taken together, these data confirm that Caf130, Btt1, Egd2, Cal4 and the N-terminus of Not1 forms an independent module with Caf130 forming the central scaffold that interacts minimally with Not1_N_, Btt1 and Cal4. The region 109-122 of Btt1 is required for its interaction with Caf130 while amino acids 1-162 of Not1 are sufficient Caf130 binding.

### Validation and structural modelling of the Caf130 module of the Saccharomyces cerevisiae CCR4-NOT complex

To validate that Caf130, Btt1, Egd2, Cal4, and residues 1-162 of Not1 (Not1_N_) form an independent structural module, we constructed operons to express these proteins recombinantly. Two plasmids were made either containing or excluding Cal4 (Figure 3A). In both cases, the Caf130 coding sequence was codon optimized, a STREP tag was appended to the N-terminus of Btt1, and a 6 His tag at the N-terminus of Caf130. Proteins were purified from extracts derived from *E. coli* grown in auto-induction media by two successive steps of affinity chromatography. Proteins in the tetrameric complex (Caf130-Btt1-Egd2-Not1_N_) co-purify efficiently and stoichiometrically (Figure 3B). While a previous study suggested that Egd2 is not part of the Caf130 module, our TAP purification data and two-hybrid assay data with a yeast strain lacking Egd2, along with this result, confirms its presence within the module. When the Cal4 coding sequence was present, an additional band corresponding to the latter protein was detected (Figure 3B). The identity of all bands was confirmed by mass spectrometry analyses (Figure S4). Taken together, these observations demonstrate that Caf130, Btt1, Egd2, Not1_N_ and Cal4 interact directly and form a stable and soluble assembly. No other yeast protein appears to be required for the formation of this complex. In particular, Cal4 is not required for the association of the other partners in a stable entity.

**Figure 3:**
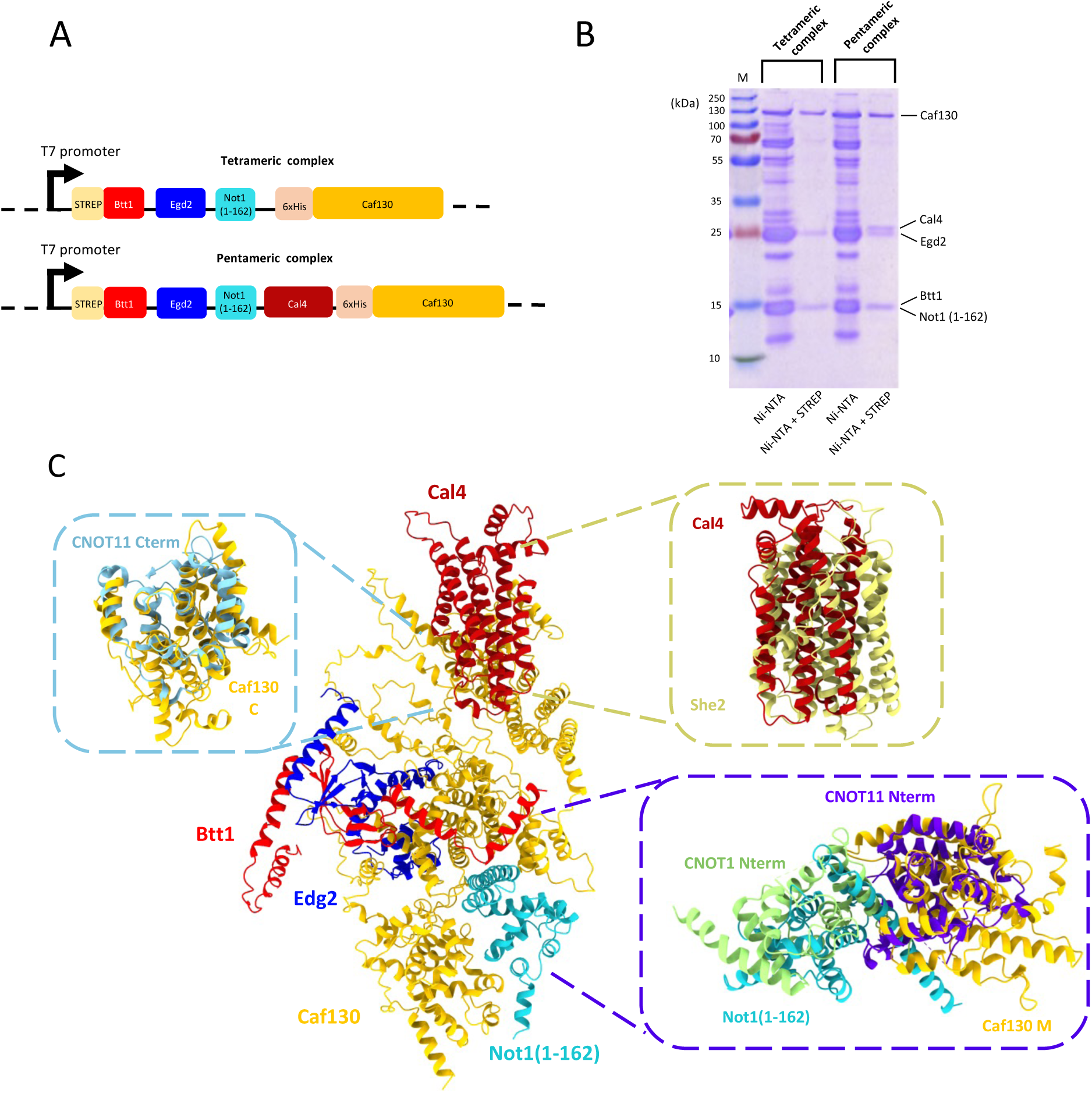
Assaying interactions between Caf130 and its partners for a structural study. (A) Schematic representation of the organization of the two operons used for recombinant protein expression. The tetrameric complex consists of STREP-Btt1, Egd2, Not1(1-162) and codon-optimized 6xHis-Caf130, while the pentameric complex includes the same factors with Cal4 in addition. (B) Coomassie-stained SDS-PAGE analysis of purifications of the tetrameric and pentameric complexes by two step affinity chromatography: first using Ni-NTA and subsequently STREP beads. Note that Egd2 (19 kDa) migrates aberrantly around 25 kDa. (C) AlphaFold3 prediction of the Caf130 module of yeast CCR4-NOT complex: Caf130 (orange), Not1_N_ (cyan), Btt1 (red), Egd2 (dark blue) and Cal4 (dark red). Top left inset: The C-terminal domain of CNOT11 (330-510) (PDB : 8BFI, (Mauxion et al. 2022)) in light blue aligns with the C-terminal region of Caf130 (694-1021) (C) in orange, with an Root Mean Square Deviation (RMSD) on CA positions (or on all atom positions) of 0.71 Å. Top right inset: The predicted structure of Cal4 (dark red) shows alignment with the determined structure of an RNA-binding protein She2 (yellow) She2 (yellow) (PDB: 1XLY, (Niessing et al. 2004)), with an RMSD of 1.16 Å. Bottom right inset: The N-terminal domain of CNOT11 (56-280) in dark blue aligns with the middle region (M) of Caf130 (298-599) in yellow, along with the 1-162 region of CNOT1_N_ (green) and Not1_N_ (cyan), with an RMSD of 1.17 Å.

As Caf130, Btt1, Egd2, Not1_N_ and Cal4 form a stable complex, we used AlphaFold3 (Abramson et al. 2024) to model its structure (Figure 3C). Consistent results were obtained and with significant scores for both intra-subunit and inter-subunit folds (Figure S5). The central core of the assembly was predicted with very high to high scores and only a limited number of peripheral elements had low or very low confidence. Caf130 appears nearly exclusively constituted of α helices and essentially divided into 3 major domains: N, M and C, along the protein N to C axis. Not1_N_ forms a globular domain that appears sandwiched between the N and M domains of Caf130 while Btt1-Egd2 adopt the previously known NAC complex structure. Both Btt1 and Egd2 contact the Caf130 M domain without interacting with Not1_N_. Interestingly, in this orientation, the residues of Btt1 identified to be required and sufficient for association with Caf130 contact directly the latter. Finally, Cal4 interacts with the Caf130 C domain without contacting other factors. Overall, this model is fully compatible with our TAP purification, two-hybrid interaction, and recombinant protein expression data.

Individual protein domains were used to find similar proteins in structural databases (Holm et Laakso 2016) while sensitive sequence similarity searches using Hidden Markov Model (HMM) (Eddy 1998) were performed to find related factors. This revealed that the Caf130 C domain sequence and fold resemble the recently described CNOT11 C-terminal domain (Mauxion et al. 2022) (Figure 3C). Moreover, the Caf130 M domain and associated Not1_N_ were structurally similar to the human CNOT11 N-terminal domain and CNOT1 N-terminal domain, respectively (Figure 3C). Together, this indicates that yeast Caf130 and human CNOT11 are related, the yeast protein lacking the extended middle region of the human factor that associates with CNOT10 and containing an additional N-terminal domain without obvious counterpart in sequence or structural databases. In addition, comparison of the predicted structure of Cal4 revealed that it resembles the yeast She2 RNA binding protein (Figure 3C-S5). Overall, these observations reveal an unexpected evolutionary link between the Caf130 module of the yeast CCR4-NOT complex and the CNOT10-CNOT11 module of its metazoan counterpart.

Plotting the evolutionary conservation of Caf130 on the predicted structure provides important information. In particular, contact sites between the different partners are usually conserved (Figure 4A-D). Intriguingly, a binding groove that accommodates an α helix of Egd2 (Figure 4D) appears more conserved than the surface discriminating Btt1 from Egd1 (Figure 4A). This may suggest that Egd2 binding is the main contributor for the NAC^Btt1^ complex binding to Caf130 while Btt1 would provide additional interaction energy as well as specificity. A few other Caf130 surfaces are also conserved and may bind transiently to other factors. Surprisingly, this includes an extensive area between the two sub-modules composing the complex (Figure 4E).

**Figure 4:**
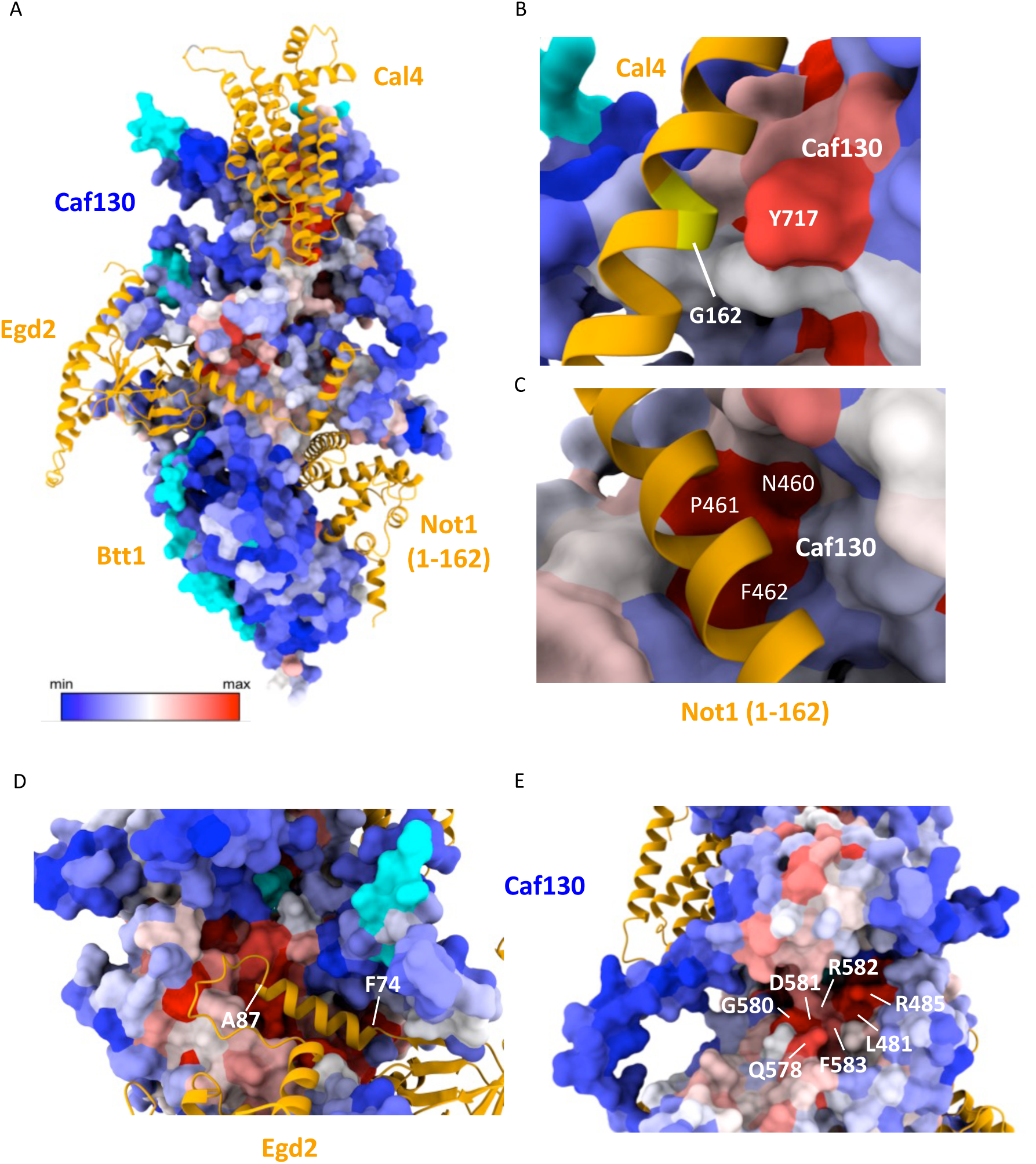
Visualization of residue conservation across different yeast species within the Caf130 module and its interacting partners. (A) The AlphaFold3 prediction of Caf130 module, annotated with residue conservation. A color gradient from the most conserved residue (red) to the least conserved (blue) represents conservation levels. The interacting partners of Caf130: Cal4, Btt1, Egd2, and Not1(1-162) are shown in orange. (B) The highly conserved residue Y717 of Caf130 (in red) is positioned opposite G162 (in yellow) on Cal4 (in orange), informing the choice of the G162V point mutation. (C) A conserved patch of Caf130, comprising residues N460, P461, and F462, is located near a helix of Not1(1-162) (in orange), supporting the selection of the point mutants N460A and P461A. (D) The second helix of Egd2 (in orange), spanning residues F74 to A87, fits into a highly conserved cavity of Caf130 (in light and dark red). (E) A highly conserved pocket in Caf130 (in light and dark red), composed of residues L481, R485, Q578, D580, D581, R582, and F583, may serve as a potential interaction region.

### Validating the AlphaFold predicted model of the Caf130 module

In the AlphaFold model, domains N and M of Caf130 contacts Not1_N_ and Egd2-Btt1 while the C domain binds Cal4 (Figure 3C). To validate this prediction, we tested using the two-hybrid assay whether Caf130 can be subdivided in domains interacting with Not1_N_ and Cal4. We observed indeed that constructs Caf130(1-622), Caf130(1-688) and Caf130(1-726) bind to Not1(1-622) while Caf130(689-1016) and Caf130 (623-1016) bind Cal4 (Figure 5A). Consistently, two-hybrid assays performed in the Δ*egd2* host strain indicate that Not1_N_ binds Caf130(1-622) or Caf130(1-688) but not Caf130(689-1016) and Caf130 (623-1016) while the reciprocal interaction was observed for Cal4 (Figure 2E).

**Figure 5:**
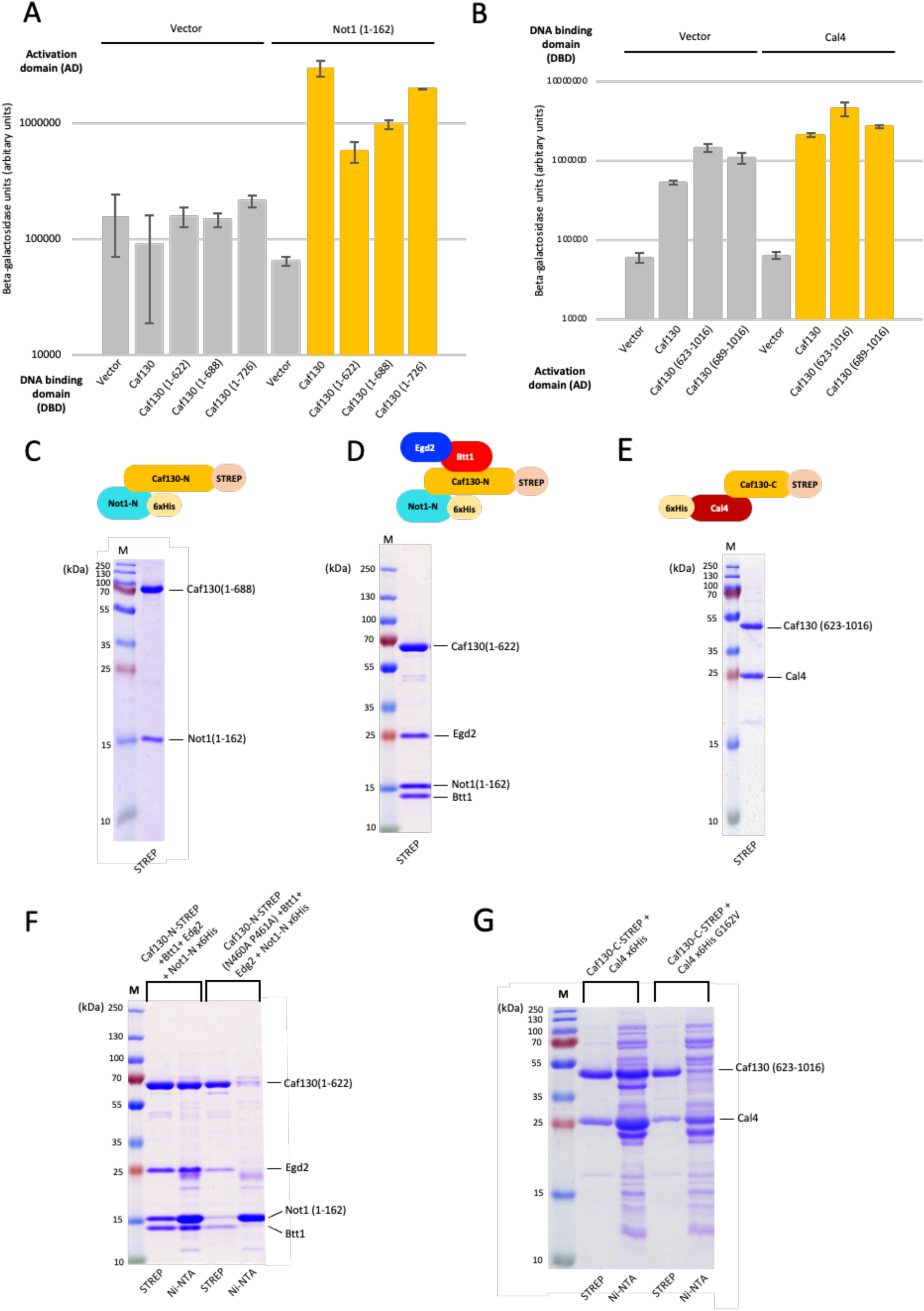
Characterization of interaction domains between Caf130 and its binding partners. (A) Two-hybrid assays indicate that the N-terminal region of Caf130 binds Not1_N_. Conditions were the same as in Fig. 2A. Positive signals were detected for Caf130, Caf130(1-622), Caf130(1-688) and Caf130 (1-726) (orange). (B) Two-hybrid assays indicate that the C-terminal region of Caf130 binds Cal4. Conditions were the same as in Fig. 2A. Positive signals were detected for Caf130, Caf130(623-1016) and Caf130 (689-1016) (orange). (C-D-E) Purification of recombinant sub-modules. At the top, schematic representations of the proteins expressed from each operon. Bottom, Coomassie-stained SDS-PAGE analyses of the single step affinity purified fractions. (C) STREP purification of Caf130 (1-688)-STREP and 6His-Not1_N_. (D) STREP purification of Caf130(1-688)-STREP, Btt1, Egd2 and 6His-Not1_N_. (E) STREP purification of Caf130(623-1016)-STREP and 6His-Cal4. (F-G) Purification of submodules carrying mutations in predicted interaction surfaces. Coomassie-stained SDS-PAGE analysis of the fraction recovered from independent Ni-NTA or STREP affinity purifications. (F) Comparison of profiles of fractions purified from operons encoding 6His-Not1(1-162)-Btt1-Egd2-Caf130(1-688)-STREP with either wild-type Caf130 or Caf130(N460A P461A). (G) Purifications of proteins expressed from an operon encoding 6His-Cal4(G162V) and Caf130 (623-1016)-STREP. Results obtained with the wild-type operon are shown as control.

To further strengthen the AlphaFold model, we constructed operons to express and purify the predicted sub-modules. A single-step Strep purification of proteins expressed from an operon encoding Caf130(1-688)-Strep and 6His-Not1_N_ resulted in a stoichiometric heterodimeric complex (Figure 5C). When the coding sequences of Btt1 and Egd2 were added to this first construct, a complex containing the four proteins Caf130(1-688)-Strep, Btt1, Egd2 and 6His-Not1_N_ was purified (Figure 5D). These results validate the existence of the corresponding sub-module and show further that Not1_N_ binds Caf130(1-688) independently of the presence or absence of the NAC. Similarly, a single-step Strep purification of proteins expressed from a plasmid encoding Caf130(623-1016) and Cal4 recovered a stoichiometric heterodimeric complex (Figure 5E). Overall, the two-hybrid and recombinant proteins expression data support the AlphaFold predicted model including regions of interaction and independent binding of the different subunits.

To get more precise validation of the interaction sites predicted by AlphaFold, we tested whether amino acid substitutions in the predicted interaction areas reduce subunit association. A highly conserved motif (N460, P461, F462) was identified in the N-terminal region of both CNOT11 and Caf130, forming a contact interface with CNOT1_N_ and Not1_N_ (Figure 4C). A double mutant N460A and P461A was introduced into the operon encoding the corresponding sub-module, 6His-Not1_N_-Btt1-Egd2-Caf130(1-688)-Strep and the resulting complex was purified by affinity chromatography using either Ni-NTA or STREP beads. Compared to the purifications of the wild-type construct that yielded roughly homogenous heterotetrameric complexes, purifications from the mutant construct with STREP beads recovered the mutated Caf130(1-688) with only reduced levels of Btt1, Egd2 and traces of Not1_N_ (Figure 5F). Conversely, affinity purification with Ni-NTA media recovered essentially Not1_N_ with only traces of the three other subunits. Altogether, these observations indicate that N460 and P461 of Caf130 are critical for Not1 binding. Further, reduced Not1 binding also decreased the co-purification of Btt1 and Egd2 suggesting that these factors may bind cooperatively possibly owing to the stabilization of the Caf130 M domain. To alter the Caf130-Cal4 interface, we substituted G162 in Cal4 with a valine in an operon encoding 6His-Cal4 and Caf130(623-1016)-Strep. The G162V was previously identified as inhibiting Cal4 function (Pillet et al. 2022). Proteins expressed from this construct and a wild-type control were purified either by Ni-NTA or STREP affinity chromatography. While heterodimeric complexes were recovered from cells containing the wild-type plasmid, proteins expressed from the mutant construct were enriched in the captured protein relative to its partner (Figure 5G) consistent with the G162V substitution impairing Cal4-Caf130 association. Taken together, the data obtained are fully consistent with the interaction scheme predicted in the structural AlphaFold model.

### Docking of the Caf130 module to ribosomes

The availability of a structural model of the Caf130 module containing the NAC^Btt1^ complex allowed us to test how the module would dock to ribosomes. Superposing the conserved NAC complex core of the AlphaFold model onto the structure of a human ribosome bound to the NAC complex (Gamerdinger et al. 2023) reveals that the Caf130 module would position near the ribosome tunnel from which emerges the nascent protein (Fig. 6A). The Caf130 module covers a small surface area of the ribosome with limited steric clashes supporting the value of this model and further arguing that the positioning of the NAC^Btt1^ on Caf130 is relevant. It also suggests that Cal4 might interact with ribosomal RNA and/or proteins (Fig. 6A).

**Figure 6:**
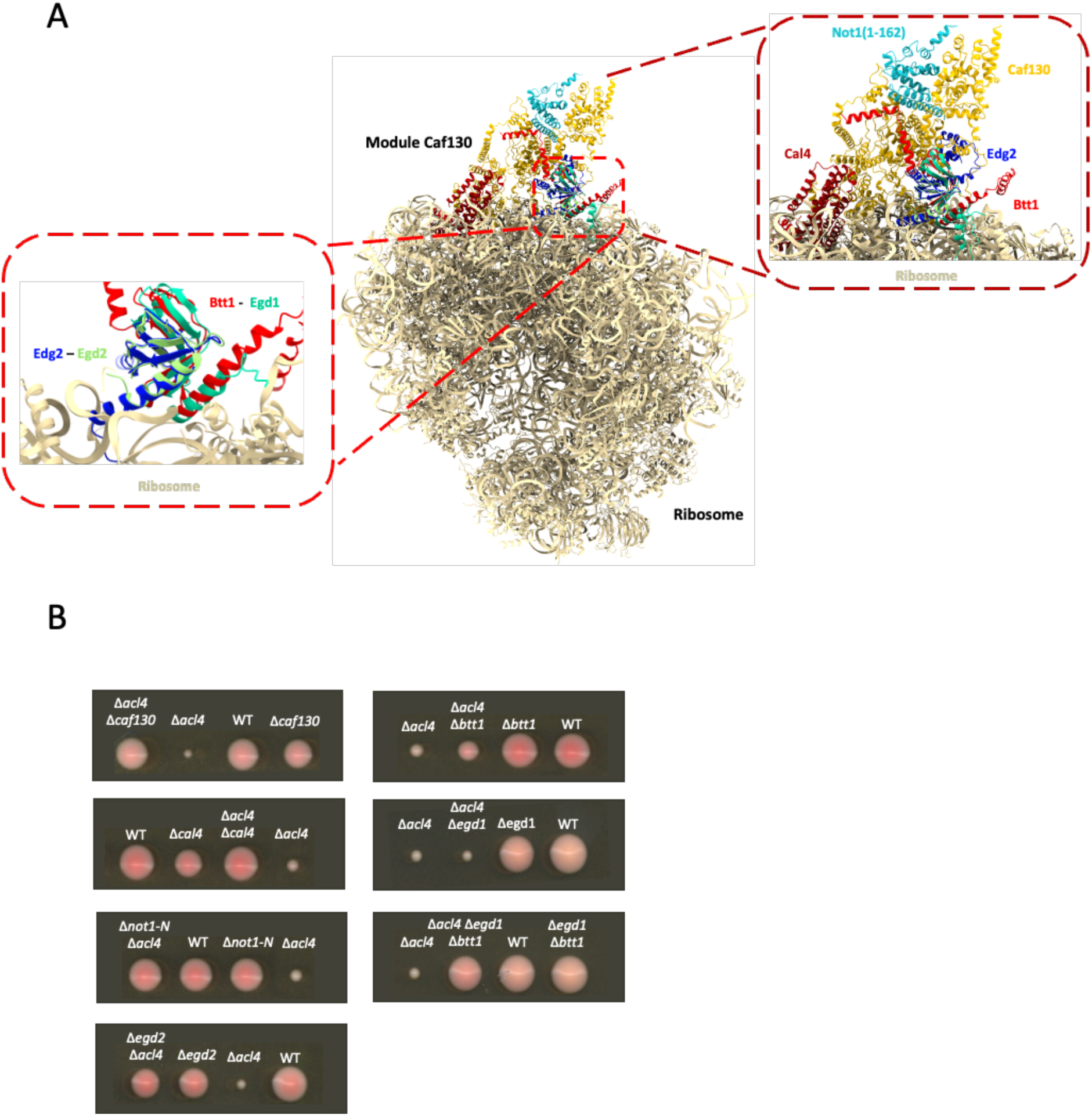
Recruitment of the CCR4-NOT N-terminal module to the ribosome is maintained by Egd1 in the absence of Btt1. (A) Docking of the Caf130 module, predicted by AlphaFold3, onto the experimentally determined ribosome structure associated to the human NAC complex (PDB : 8P2K, (Gamerdinger et al. 2023)).The ribosome is depicted in beige, with the human NAC complex (Egd1 (green) and Egd2 (light green). The Caf130 module is visualized with Caf130 (orange), Btt1 (red), Egd2 (dark blue), Not1N (cyan) and Cal4 (dark red). (B) Suppression of the Δ*acl4* mutation by mutants of the Caf130 module. Growth was monitored by following the size of colonies resulting from spored issues from the sporulation of heterozygous diploid cells on YPDA plates. Plate images were captured 5 days post-dissection.

### Assaying the Caf130 module function reveals overlapping roles of Btt1 and Egd1

Functional analyses were performed using the suppression of the growth defect conferred by acl4 inactivation that occurs when Caf130 is inactivated (Pillet et al. 2022). Consistent with the previous results, deletions of *caf130*, *cal4* or *not1_N_* almost completely suppressed the *Δacl4* growth defect while having themselves little impact on growth (Figure 6B). Even though Egd2 was reported not to be associated with the Caf130 module (Pillet et al. 2022), its deletion also rescued the *Δacl4* growth defect (Figure 6B). This supports its implication in the same biological pathway as its partners and strongly argues for its presence in the Caf130 module, in agreement with affinity purification results and AlphaFold modeling. While inactivation of Egd1 had no suppressing effect as anticipated, deletion of *btt1* resulted in surprisingly only a partial suppression (Figure 6B). However, full suppression was detected when both Btt1 and Egd1 were inactivated (Figure 6B). This observation reveals overlapping roles of Btt1 and Egd1. This suggests that, in the absence of *btt1*, Caf130 can interact with the Egd1/Egd2 NAC complex albeit more weakly than with the Btt1/Egd2 complex. This interaction would still enable the CCR4-NOT complex to associate with ribosomes translating Rpl4, particularly when Rpl4 encounters folding difficulties. Several observations support this proposal including the strong similarity between the NAC^Btt1^ and NAC^Egd1^ complexes that would both associate to Caf130; the main contacts being provided by Egd2 while Egd1 or Btt1 would only modulate the final binding affinity.

## Discussion

Large-scale proteomic screens (Ito et al. 2001; Krogan et al. 2006; Cui et al. 2008) and interaction analyses conducted within our team revealed an association between Caf130, a subunit of the CCR4-NOT complex, and Btt1, a β-subunit of the NAC complex. Subsequent suppressor screens for the slow-growth phenotype resulting from ACL4 inactivation in yeast (Pillet et al. 2022b) further supported this interaction. These observations established an unexpected physical link between translation—specifically the exit of the nascent peptide from the ribosome—and mRNA degradation, initiated by deadenylation. However, the organization and function of all factors involved in this mechanism remain poorly understood. To extend these preliminary findings, we initially set out to identify Caf130’s interaction partners by purifying the protein using Tandem Affinity Purification (TAP), followed by mass spectrometry identification of co-purified proteins. This approach detected both expected and unexpected partners (Figure 1A). Nearly all subunits of the CCR4-NOT complex were recovered (Not1, Not3, Not4, Not5, Caf130, Caf40, CCR4, and Caf1), with the exception of Not2. Since Not2 was also undetected in the extract—likely due to its small size—its absence is not informative. Not3, Not4, and Not5 were detected but at lower levels than other subunits, suggesting potential loss during purification. Supporting this, a previous study reported dissociation of the CCR4-NOT complex during the calmodulin purification step (Nasertorabi et al. 2011). In this case, as these subunits interact at the C-terminus of Not1— far from the site where Not1 binds Caf130—their loss is more likely due to limited proteolysis or partial dissociation during purification.

The yeast two-hybrid system confirmed the interaction between Caf130 and Not1 (Figure 2A). While this does not demonstrate a direct interaction, it suggests that Caf130 either binds Not1 directly or interacts with another CCR4-NOT subunit. Analyses using a truncated version of Not1 revealed that amino acids 1–162 are sufficient for binding to Caf130 (Figure 2D). The strong positive signal detected with Not1(1–162) is particularly informative, as this region is relatively small and not known to bind other CCR4-NOT subunits, reinforcing the hypothesis of a direct Caf130-Not1 interaction (Caulier et al., 2025). My biochemical studies with recombinant proteins, along with the AlphaFold model, confirmed this direct interaction (Figures 3, 5). Additionally, another yeast two-hybrid analysis, mapping Caf130’s interaction regions with its partners, showed that Not1(1–162) interacts with the N-terminal region of Caf130, which can be reduced to amino acids 1–622 (Figure 5C). These results provide further detail on the interaction between these proteins. Furthermore, aligning the sequences of the N-terminal regions from the experimental structures of human CNOT11 and CNOT1 (Mauxion et al. 2022) with those of yeast Caf130 and Not1 from our AlphaFold model revealed a similarity in interactions between the human and yeast CCR4-NOT complexes (Figure 3C). This conservation is also evident in a conserved region between the C-terminus of CNOT11 and another region of Caf130 (Figure 3C). These observations suggest that the N-terminal regions of the human and yeast CCR4-NOT complexes share a common evolutionary origin and raise the question of their functional similarity.

The C-terminal region of CNOT11 has been described as acting as an “antenna,” facilitating interactions between the N-terminal region of the CCR4-NOT complex and numerous partners, including GGNBP2 (Mauxion et al. 2022) and SCAPER, the latter of which is involved in a quality control mechanism similar to that of Caf130 (Höpfler et al. 2023b).

The Caf130 interactome revealed a novel partner of unknown function, Cal4 (Figure 1A). Independently, Cal4 was identified in a suppressor screen for the slow-growth phenotype resulting from ACL4 inactivation (Pillet et al. 2022b). A yeast strain expressing TAP-tagged Cal4 was constructed, allowing the identification of its partners after TAP purification. Cal4’s interactome proved largely similar to that of Caf130 (data not shown). Yeast two-hybrid results confirmed the interaction between Caf130 and Cal4 and showed that the C-terminal region of Caf130, reducible to amino acids 689–1016, interacts with Cal4 (Figures 2 and 5). Co-purification of Caf130(623–1016) and Cal4 confirmed that this interaction is direct (Figure 5E). These data confirm that Cal4 is a bona fide Caf130 partner and highlight the conserved role of the CNOT11 C-terminal domain in partner recruitment. Thus, Cal4 appears to be a new subunit of the CCR4-NOT complex. Notably, a highly sensitive sequence similarity analysis (HMM) revealed a similarity between Cal4 and an RNA-binding protein named She2. This similarity was confirmed by aligning the experimental structure of She2 with the AlphaFold-generated model of Cal4 (Figure 3C). This could suggest that Cal4 also has the ability to bind RNA.

Our proteomic analyses confirmed the presence of the NAC(Btt1) complex in Caf130’s interactome while suggesting that the NAC(Egd1) complex is underrepresented (Figure 1A). Given that Egd1 is 20-fold more abundant than Btt1 (Breker, Gymrek, and Schuldiner 2013; Dénervaud et al. 2013), this conclusion is reinforced. Yeast two-hybrid results confirmed that Caf130’s interaction with Btt1 is specific, with at best a weak signal detected for Egd1 (Figure 2). This differential signal, combined with the high similarity between Btt1 and Egd1,

facilitated the creation of chimeras, which allowed mapping of a critical Btt1 region required for binding to Caf130. Yeast two-hybrid tests with truncations and chimeras identified Btt1’s 109–122 region as necessary and sufficient for interaction with Caf130 in the context of a β-NAC scaffold that enables association with NAC α (Figures 2 and S2).

Yeast two-hybrid assays did not produce signals suggesting an interaction between Egd2 and Caf130. Given the observed signal between Btt1 and Caf130 in yeast two-hybrid assays, TAP purification results, recombinant complex production, and structural modeling, one might have expected a positive signal between Caf130 and Egd2 as well. However, yeast two-hybrid tests can yield negative results for many reasons. For instance, Egd2-containing fusions may be poorly expressed, inefficiently transported to the nucleus, block transcriptional activation, or be titrated out by Egd1. In conclusion, even though yeast two-hybrid assays do not reveal an interaction between Egd2 and Caf130, our other data strongly support this association.

The authors of the study on suppressors of the slow-growth phenotype conferred by the acl4 mutation questioned the presence of Egd2 within the Caf130 module (Pillet et al. 2022b). To address this, we validated the direct interaction by reconstituting the entire module in an operon: Btt1(STREP)-Egd2-Not1N-Cal4-Caf130(6xHis), expressed in a bacterial system, followed by dual purification. This method allowed us to demonstrate a direct interaction among all the module’s proteins, eliminating potential cofactors present in yeast that could interfere with yeast two-hybrid analyses (Figure 3B). The results showed that all proteins, including Egd2, co-purified efficiently and stoichiometrically. Protein identities were confirmed by mass spectrometry (Figure S4). Combined with Caf130 interactome data (Figure 1) and yeast two-hybrid data in a yeast strain lacking egd2 (Figure 2E), these results allow us to affirm that Egd2 is indeed part of the module.

This purification method, along with yeast two-hybrid data and our AlphaFold model, also enabled more precise mapping of Caf130’s interaction region with the NAC complex. Specifically, co-purification of the Caf130(1–622)(STREP)-Btt1-Egd2-Not1N(6xHis) complex revealed that residues 1–622 of Caf130 are sufficient for interaction with the NAC complex and that this interaction is enhanced in the presence of Not1N (Figure 5D). Additionally, our mass spectrometry results with the Caf130-TAP Not1 (ΔN-terminal) strain revealed a decreased association of Btt1 and Egd2 with Caf130 compared to the wild-type Caf130-TAP strain (Figure 1B). These results argue for a role of Not1’s N-terminal region in promoting or stabilizing Caf130’s interaction with the NAC complex.

Initial structural modeling of the complete module was performed using AlphaFold2 (data not shown). The 25 predictions obtained were superimposed, revealing that the relative positions of Cal4 and Not1N were consistent within the complex. However, two distinct orientations were observed for Btt1 and Egd2. In the first, the helix corresponding to Btt1’s 109–122 region (required for interaction with Caf130) was in contact with Caf130, while in the second orientation, this helix pointed outward. We concluded that the first orientation was correct, allowing us to more precisely define the contact region between Caf130 and Btt1. We repeated our predictions with AlphaFold3, which now only predicts the correct orientation. The accuracy of these predictions was validated by yeast two-hybrid data and recombinant complex analysis (Figure 5). AlphaFold represents a revolutionary advance in protein structure prediction (Jumper et al. 2021). But, while AlphaFold is a powerful tool, it must be used in conjunction with other experimental approaches to confirm the predictions.

To better identify interaction regions and confirm our model, point mutants were generated in the two entities constituting the Caf130 module. For the entity containing Caf130’s C-terminal region and Cal4, mutants were obtained during the analysis of acl4 suppressors (Pillet et al. 2022b). These mutants were primarily located in the interaction region between these two proteins or likely affected their folding. Other surfaces were unaffected. Point mutations were introduced into plasmids containing the coding sequence for Caf130’s C-terminal region (R712G and E836K), as well as in Cal4 (G162V and L208P), and the corresponding recombinant proteins were purified. Among the tested mutations, only Cal4(G162V) significantly impacted its co-purification with Caf130’s C-terminal domain, consistent with its positioning opposite a conserved region of Caf130 in the predicted complex structure (Figures 4B and 5G). The other mutations, R712G and E836K in Caf130 and L208P in Cal4, did not affect the interaction of the recombinant proteins, although we cannot rule out changes in affinity or other in vivo parameters. In some cases, the mutated protein was no longer expressed, suggesting an effect on protein stability (data not shown). To finalize validation of this interaction region, the association between separately purified Caf130C and Cal4, or Cal4(G162V), could be tested using ITC and/or Nano-DSF.

For the Caf130N-Not1N-Btt1-Egd2 entity, evolutionary conservation analysis revealed that three amino acids—N460, P461, and F462—in Caf130’s N-terminal region form an interaction interface with Not1, and this interface is conserved in the human CNOT11-CNOT1 complex (Figure 4C). Consequently, we mutated two of these residues, N460A and P461A, in the plasmid encoding Caf130N-Not1N-Btt1-Egd2 and purified the resulting recombinant proteins. The results showed that these mutations significantly reduced the interaction between Caf130 and Not1, validating the interaction region (Figure 5F). Furthermore, this reduction in Not1-Caf130 interaction was accompanied by a decrease in Caf130-NAC interaction (Figure 5F), consistent with the effect of deleting Not1’s N-terminal region, which also reduces NAC-Caf130 interaction in vivo (Figure 1B).

In addition to the CCR4-NOT and NAC(Btt1) complexes, numerous ribosomal proteins were identified in Caf130’s interactome (Figure 1A). The absence of some low-molecular-weight ribosomal proteins—rich in arginine and lysine and thus digested into smaller fragments by trypsin during sample preparation—can be explained by identification issues, as these proteins generate few identifiable peptides. Given this, it is likely that a complete ribosome co-purified with Caf130. The enrichment levels of ribosomes in the fraction purified with Caf130 were not very high, but this may result from Caf130 being associated with only a subset of ribosomes. To validate Caf130’s specific association with ribosomes, we used a direct approach with polysome gradients. This observation was confirmed by analyzing Cal4’s association with polysomes (Appendix 1). Overall, the collected information supports an interaction between the Caf130 module and ribosomes. A model of this interaction can be proposed by superimposing our AlphaFold-generated model onto a ribosome structure associated with NAC (Figure 6A). This superposition was possible due to the strong similarity between human and yeast NAC, emphasized by the conservation of two α-helices—one from Egd1 (and Btt1 in yeast) and the other from Egd2. These helices are oriented toward the ribosome, enabling interaction at the tunnel exit. In this configuration, the Caf130 module is precisely positioned at the peptide exit tunnel of the ribosome, ready to interact with nascent peptides. This positioning strongly supports the initially described quality control model, where Btt1 binds to the Rpl4 protein as it emerges from the ribosomal tunnel and recruits the CCR4-NOT complex to degrade the corresponding mRNA when its chaperone is absent or insufficient. It is also noteworthy that in this model, Cal4 is located near the 28S ribosomal RNA. As previously mentioned, a similarity between Cal4 and She2, an RNA-binding protein, was detected. This suggests that Cal4 might potentially interact with 28S rRNA. Techniques such as gel-shift assays, RIP-PCR, or CLIP-seq could be used to test this hypothesis.

The detection of the Pbp1 complex (Pbp1, Mkt1, Pbp4, and Lsm12) among the proteins associated with Caf130 is also interesting (Figure 1A), as it contributes to modulating poly(A) tail length and translation regulation (Mangus, Amrani, and Jacobson 1998). Further analyses will be necessary to determine whether this interaction has a biological role or if the co-purification merely results from random associations of these factors with the same mRNAs.

To explore the biological and physiological consequences of the various factors, we adapted a phenotypic test: the functionality of the Caf130 module was evaluated by measuring its impact on the growth of yeast lacking Acl4, the specific chaperone of Rpl4. Deleting the gene encoding ACL4 significantly reduces yeast vegetative growth. However, this growth defect is suppressed when the CAF130 gene is also deleted (Pillet et al. 2022b). This suppression occurs because, in the simultaneous absence of Acl4 and Caf130, the mRNA levels of RPL4 increase, leading to enhanced synthesis of this protein. The lack of CCR4-NOT complex recruitment when Caf130 is absent results in this increase (likely due to reduced deadenylation), which in turn leads to increased synthesis of the ribosomal protein Rpl4 and, consequently, better production of the 60S ribosomal subunit, thus promoting better cell growth.

This strategy allowed us to evaluate the phenotypes of the different partners in the Caf130 module. Our results show that the absence of Caf130, Cal4, Not1N, or Egd2 leads to near-complete suppression of the growth phenotype associated with the lack of Acl4 (Figure 6B). In contrast, we observed that the absence of Btt1 only partially suppresses this phenotype, suggesting that Btt1 has a limited role in the mechanism. Deleting its paralog, EGD1, does not affect the growth phenotype associated with the absence of Acl4. However, the combined deletions of btt1 and egd1 fully suppress the growth phenotype. This observation suggests that, in the absence of Btt1, Caf130 can interact with Egd1-Egd2, albeit more weakly than with Btt1-Egd2, thereby maintaining CCR4-NOT complex recruitment via Caf130 to ribosomes translating Rpl4 when its folding is slow. This result supports the hypothesis of an interaction between Caf130 and Egd1. Yeast two-hybrid results are also compatible with this model. Thus, these data explain why egd2 deletion causes complete suppression of the phenotype: its absence prevents Caf130 from binding to both Btt1 and Egd1, thereby blocking CCR4-NOT complex recruitment to ribosomes translating the RPL4 protein).

Thus, the coupling of translation and degradation benefits cells by enabling rapid regulation of the quantity of certain proteins, especially under stress or malfunction conditions. This mechanism is essential for managing misfolded and/or potentially harmful proteins that can disrupt cellular function. By eliminating misfolded proteins, cells prevent their accumulation, thereby reducing the risk of aggregate formation and ensuring quality control while conserving resources by preventing their useless accumulation.

Interestingly, mapping the sequence conservation of Caf130 onto its AlphaFold-predicted structure revealed a highly conserved region (Figure 4D). This region forms a groove into which an Egd2 helix inserts, strongly suggesting a direct and important contact between these two proteins. This observation implies that Btt1 merely increases NAC’s affinity for Caf130 compared to the affinity of NAC composed of Egd2 and Egd1. This aligns with the suppression phenotypes observed for Δegd2, Δbtt1, Δegd1, and Δbtt1Δegd1 (Figure 6B). It would be interesting to generate point mutants in both Caf130’s conserved region and the adjacent Egd2 helix to test whether the interaction between Caf130 and NAC is still maintained. Once validated by interaction tests, these mutants could be functionally validated using the phenotypic test, following the same principle as for previous mutants (Caf130 N460, P461, and Cal4 G162V).

Our data, along with published findings, corroborate the importance of the CCR4-NOT complex not only for its deadenylase activity but also for the functions of its various subunits (Caulier et al., 2025). These subunits confer an essential role in diverse cellular processes, particularly in the co-translational regulation of certain ribosomal proteins, as described in this study. The CCR4-NOT complex can thus adapt to the specific needs of cells by directing its activity or interactions to maintain cellular process balance.

## Materials and methods

### Yeast strains, genetic methods and plasmids

All strains used in this study are derived from the W303 background, except for the strain employed for the yeast two-hybrid assays, which was derived from MAV203 (Invitrogen). Chromosomal gene deletions were carried out using PCR fragments, following standard methods (*Guthrie C et al., 2002)*. The strain expressing TAP-tagged Caf130 was constructed as described previously (Rigaut *et al*., 1999). Combinations of mutations or genetic variants were generated by crossing, followed by sporulation of the resulting diploids, tetrad dissection and validation of haploid spore clones with the desired genotype. The Caf130-TAP Not1Δ*N(1-162)* strain was generated using a « pop-in/pop-out » strategy with the *URA3* selectable marker and 5-FOA counterselection (*Guthrie C et al., 2002)*. Recombinant DNA manipulations were conducted following standard molecular biology protocols. *E. coli* MH1 was used for transformation and plasmid amplification. All DNA fragments generated by PCR amplification were verified by sequencing.

### TAP-tagged protein purification

TAP purification was performed as described by Rigaut et al. (1999). Briefly, TAP-tagged strains were grown in 2 L of YPDA medium to an OD₆₀₀ of 2. Cells were harvested, washed, and lysed using a Constant Systems disruptor in buffer A. The lysate was adjusted to 0.2 M KCl, clarified by ultracentrifugation, and dialyzed in buffer D. Protein extracts were loaded onto IgG agarose beads (Sigma), and after washing, TEV protease cleavage was performed at 16 °C. Eluates were applied to a calmodulin affinity resin (Stratagene), and bound complexes were eluted in EGTA-containing buffer. Fractions 2 to 5, confirmed by Western blot to contain Caf130, were pooled, TCA-precipitated, and analyzed by mass spectrometry. Detailed buffer compositions are available in Supplementary information.

### Yeast Two-Hybrid Assay

Protein–protein interactions were tested using the ProQuest yeast two-hybrid system (Thermo Fisher Scientific). Plasmids encoding the proteins of interest were co-transformed into the MAV203 yeast strain, and transformants were selected on SC medium lacking leucine and tryptophan (-Leu, -Trp). For each assay, three independent yeast cultures were seeded from distinct colonies (three transformants per culture) and grown in 3 mL -Leu -Trp selective media at 30°C for 18 h. Cultures were subsequently diluted tenfold, and O.D600 was measured using microplate reader (SYNERGY HTX). 50 μL of diluted cells were mixed with 50 μL of Beta-Glo reagent (Promega) and incubated in the dark at room temperature for 60 minutes. Luminescence was quantified using a Centro XS3 LB 960 luminometer (Berthold Technologies).

### Sequence alignments and structural analyses

Protein sequences from various organisms, including human and yeast, were retrieved from databases such as UniProt and SGD. FASTA files were compiled and aligned using Clustal W. The structure of the Caf130 module was predicted using Alphafold 2 et AlphaFold 3. Visualisation and structural alignments were performed using ChimeraX (https://www.cgl.ucsf.edu/chimerax/), based on the Caf130 predicted module and the following reference structures: human CNOT11 (residues 56–280 and 330–510) (PDB: 8BFI, Mauxion et al., 2022), yeast She2 (PDB: 1XLY, Niessing et al., 2004), and the 80S ribosome associated with the human NAC (PDB: 8P2K, Gamerdinger et al., 2023).

### Purification of recombinant proteins

The tetrameric complex (Btt1, Egd2, Not1(1–162), Caf130) and the pentameric complex (Btt1, Egd2, Not1(1–162), Cal4, Caf130) were each sub-cloned as separate operons. In both constructs, codon-optimized Caf130 was fused at its N-terminus to a 6xHis-tag, while Btt1 was tagged at its N-terminus with a STREP tag. Complexes were expressed separately in BL21 CodonPlus-RIL in Terrific Broth (TB) auto-induction medium. The pellets were resuspended in 10mM imidazole, 300mM NaCl, 0,1% NP40, PBS 1X, supplemented with 1X PIC (Roche) and 0.1mM PMSF and lysed by sonication. The proteins were purified by nickel affinity chromatography using Ni-NTA agarose beads (Qiagen) at 4 °C for 1 hour, according to the manufacturer’s instructions. Following elution, the complex was further purified by Strep affinity chromatography using Strep-Tactin XT 4Flow beads (IBA) and Buffer W (IBA) (100 mM Tris-Cl pH 8.0, 150 mM NaCl, 1 mM EDTA pH 8.0) at 4°C for 1 hour, also following the manufacturer’s protocol.

The recombinant sub-modules Caf130(1–688)-Strep with 6xHis-Not1(1–162); Caf130(1–688)-Strep with Btt1, Egd2, and 6xHis-Not1(1–162); and Caf130(623–1016)-Strep with 6xHis-Cal4 were each sub-cloned as separate operons, expressed in *E. coli* BL21 CodonPlus-RIL, and purified by only Strep affinity chromatography as described above, following the manufacturer’s instructions.

### Phenotypic assay of the Caf130 module activity within the CCR4-NOT complex

The function of the Caf130 module was assessed based on its ability to suppress the slow-growth phenotype caused by *ACL4* deletion (*Δacl4*) (Pillet et al., 2022). Several strains were generated by crossing *Δacl4* strains with mutants of various Caf130 module components (Δ*caf130,* Δ*cal4*, Δ*not1(1-162),* Δ*btt1,* Δ*egd2*, Δ*egd1,* Δ*btt1*Δ*egd1).* Sporulation and tetrad dissection were performed (*Guthrie C et al., 2002*. Colony size was monitored over 5 days, with daily imaging starting on day 3. Segregation of *Δacl4* and other gene deletions was confirmed using selection markers. For the *Not1-ΔN* allele, which lacks a selectable marker, segregation was verified by PCR using primers flanking the deleted region.

## Acknowledgements

We thank the members of Dr. Bertrand Séraphin’s laboratory for insightful comments, and helpful suggestions throughout this study. We also thank the IGBMC mass spectrometry and protein purification platforms for their technical assistance.

## Author Contributions Statement

G.C.: Experimental design, methodology, investigation, formal analysis, validation, visualization, writing – original draft, writing – review and editing.

F.M.: Conceptualization, methodology, supervision, formal analysis, validation, manuscript writing, review and editing.

B.S.: Conceptualization, methodology, investigation, formal analysis, supervision, funding acquisition, resources, formal analysis, writing of original dra, review and editing.

## Bibliography

Abramson, J., Adler, J., Dunger, J., Evans, R., Green, T., Pritzel, A., et al. (2024). Accurate structure prediction of biomolecular interactions with AlphaFold 3. Nature, 630, 493–500.

Absmeier, E., Chandrasekaran, V., O’Reilly, F. J., Stowell, J. A. W., Rappsilber, J., & Passmore, L. A. (2023). Specific recognition and ubiquitination of translating ribosomes by mammalian CCR4–NOT. Nature Structural & Molecular Biology, 30, 1314–1322.

Basquin, J., Roudko, V. V., Rode, M., Basquin, C., Séraphin, B., & Conti, E. (2012). Architecture of the nuclease module of the yeast Ccr4–Not complex: The Not1– Caf1–Ccr4 interaction. Molecular Cell, 48, 207–218.

Bhaskar, V., Roudko, V., Basquin, J., Sharma, K., Urlaub, H., Séraphin, B., et al. (2013). Structure and RNA-binding properties of the Not1–Not2–Not5 module of the yeast Ccr4–Not complex. Nature Structural & Molecular Biology, 20, 1281– 1288.

Bhaskar, V., Basquin, J., & Conti, E. (2015). Architecture of the ubiquitylation module of the yeast Ccr4–Not complex. Structure, 23, 921–928.

Buschauer, R., Matsuo, Y., Sugiyama, T., Chen, Y.-H., Alhusaini, N., Sweet, T., et al. (2020). The Ccr4–Not complex monitors the translating ribosome for codon optimality. Science, 368, eaay6912.

Caulier, G., Siblini, J., Sène, L., Mauxion, F., & Séraphin, B. (2025). The CCR4–NOT complex: A multifaceted sensor of molecular signals instructing eukaryotic mRNA translation and stability. Nucleic Acids Research, 53, gkaf1401.

Collart, M. A. (2016). The Ccr4–Not complex is a key regulator of eukaryotic gene expression. WIREs RNA, 7, 438–454.

Cui, Y., Ramnarain, D. B., Chiang, Y.-C., Ding, L.-H., McMahon, J. S., & Denis, C. L. (2008). Genome wide expression analysis of the CCR4-NOT complex indicates that it consists of three modules with the NOT module controlling SAGA-responsive genes. Molecular Genetics and Genomics, 279, 323–337.

Denis, C. L. (1984). Identification of new genes involved in the regulation of yeast alcohol dehydrogenase II. Genetics, 108, 833–844.

Eddy, S. R. (1998). Profile hidden Markov models. Bioinformatics, 14, 755–763.

Gamerdinger, M., Jia, M., Schloemer, R., Rabl, L., Jaskolowski, M., Khakzar, K. M., et al. (2023). NAC controls cotranslational N-terminal methionine excision in eukaryotes. Science, 380, 1238–1243.

Guthrie, C., & Fink, G. R. (Eds.). (2002). Guide to Yeast Genetics and Molecular and Cell Biology, Part B. Methods in Enzymology, 350. Academic Press.

Holm, L., & Laakso, L. M. (2016). Dali server update. Nucleic Acids Research, 44, W351–W355.

Ito, T., Chiba, T., Ozawa, R., Yoshida, M., Hattori, M., & Sakaki, Y. (2001). A comprehensive two-hybrid analysis to explore the yeast protein interactome. Proceedings of the National Academy of Sciences of the United States of America, 98, 4569–4574.

Krogan, N. J., Cagney, G., Yu, H., Zhong, G., Guo, X., Ignatchenko, A., et al. (2006). Global landscape of protein complexes in the yeast Saccharomyces cerevisiae. Nature, 440, 637–643.

Liu, H.-Y., Badarinarayana, V., Audino, D. C., Rappsilber, J., Mann, M., & Denis, C. L. (1998). The NOT proteins are part of the CCR4 transcriptional complex and affect gene expression both positively and negatively. The EMBO Journal, 17, 1096–1106.

Mauxion, F., Prève, B., & Séraphin, B. (2013). C2ORF29/CNOT11 and CNOT10 form a new module of the CCR4-NOT complex. RNA Biology, 10, 267–276.

Mauxion, F., Basquin, J., Ozgur, S., Rame, M., Albrecht, J., Schäfer, I., et al. (2023). The human CNOT1-CNOT10-CNOT11 complex forms a structural platform for protein-protein interactions. Cell Reports, 42, 111902.

Niessing, D., Hüttelmaier, S., Zenklusen, D., Singer, R. H., & Burley, S. K. (2004). She2p is a novel RNA binding protein with a basic helical hairpin motif. Cell, 119, 491–502.

Oberholzer, U., & Collart, M. A. (1998). Characterization of NOT5 that encodes a new component of the Not protein complex. Gene, 207, 61–69.

Pillet, B., Méndez-Godoy, A., Murat, G., Favre, S., Stumpe, M., Falquet, L., et al. (2022). Dedicated chaperones coordinate co-translational regulation of ribosomal protein production with ribosome assembly to preserve proteostasis. eLife, 11, e74255.

Puig, O., Caspary, F., Rigaut, G., Rutz, B., Bouveret, E., Bragado-Nilsson, E., et al. (2001). The tandem affinity purification (TAP) method: A general procedure of protein complex purification. Methods, 24, 218–229.

Rigaut, G., Shevchenko, A., Rutz, B., Wilm, M., Mann, M., & Séraphin, B. (1999). A generic protein purification method for protein complex characterization and proteome exploration. Nature Biotechnology, 17, 1030–1032.

Sakai, A., Chibazakura, T., Shimizu, Y., & Hishinuma, F. (1992). Molecular analysis of POP2 gene, a gene required for glucose-derepression of gene expression in Saccharomyces cerevisiae. Nucleic Acids Research, 20, 6227–6233.

Wahle, E., & Winkler, G. S. (2013). RNA decay machines: Deadenylation by the Ccr4–Not and Pan2–Pan3 complexes. Biochimica et Biophysica Acta – Gene Regulatory Mechanisms, 1829, 561–570.

Zhu, X., Cruz, V. E., Zhang, H., Erzberger, J. P., & Mendell, J. T. (2024). Specific tRNAs promote mRNA decay by recruiting the CCR4-NOT complex to translating ribosomes. Science, 386, eadq8587.

